# Enhancing a multipurpose artificial urine for culture and gene expression studies of uropathogenic *Escherichia coli* strains

**DOI:** 10.1101/2023.10.13.562160

**Authors:** Patricia T. Rimbi, Nicky O’Boyle, Gillian R. Douce, Mariagrazia Pizza, Roberto Rosini, Andrew J. Roe

## Abstract

Uropathogenic *Escherichia coli* (UPEC) are the most common cause of urinary tract infections, which pose a great burden on global health and the economy through morbidity, mortality, healthcare costs and loss of productivity. Pooled human urine can be used as a growth medium for *in vitro* studies, however even if the same donors are used, composition can vary depending largely on diet and fluid intake. There have been a number of artificial urine formulae used as alternatives to pooled human urine. However, we observed that a recently reported multipurpose artificial urine was unable to support the growth of prototypic UPEC strains suggesting it lacked key metabolites. We therefore used liquid chromatography mass spectrometry to identify and adjust the metabolic profile of multipurpose artificial urine closer to that of pooled human urine. Modification in this way facilitated growth of UPEC strains with growth rates similar to those obtained in pooled human urine. Transcriptomic analysis of UPEC strains cultured in enhanced artificial urine and pooled human urine showed that the gene expression profiles are similar, with less than 7% of genes differentially expressed between the two conditions. The data support this enhanced artificial urine as a robust media to study aspects of UPEC physiology *in vitro*.

## Introduction

Uropathogenic *Escherichia coli* (UPEC) strains are the leading cause of urinary tract infections, which have associated morbidities, mortality, and economic burden. UPEC strains disseminate from the gastrointestinal tract and colonise the periurethral area, from which they can invade and colonise the urinary tract. Hence, urine is arguably the most appropriate growth medium to culture UPEC *in vitro* and for studying gene expression, however there are many hurdles associated with the use of urine. These challenges include variations in composition (Sarigul et al., 2019), the need for donors and the stability of urine over time. Therefore, to better understand UPEC physiology *in vitro* we need a growth medium that is physiologically relevant and highly reproducible. Artificial urine (AU) has been used for different purposes in the literature, for example to examine the expression of virulence genes in UPEC strain CFT073 during cell adhesion (Sarshar et al., 2022). Certain formulations have components that are not normally found in healthy human urine, such as bicarbonate (Brooks and Keevil, 1997) and therefore are not the most accurate representation of the chemical profile of healthy human urine.

Analytical techniques such as mass spectrometry, nuclear magnetic resonance and liquid chromatography mass spectrometry have been combined to demonstrate that human urine is a complex medium, with thousands of components (Sarigul et al., 2019). Based on this analysis, a recent publication formulated an artificial urine that tried to replicate healthy human urine (Sarigul et al., 2019). The authors compared their multi-purpose artificial urine (MP-AU) to urine collected from 28 individuals by using attenuated total reflection-Fourier transform infrared spectroscopy (Sarigul et al., 2019). The 13 components of MP-AU were shown to be at more physiologically relevant levels compared to the two other artificial urine formulations (Brooks and Keevil, 1997; Chutipongtanate and Thongboonkerd, 2010; Sarigul et al., 2019). In this study, we aimed to determine whether Sarigul’s MP-AU could be used as an alternative to human urine in *in vitro* studies of UPEC strains. We have taken the pre-existing MP-AU and enhanced it with specific metabolites present in human urine to support the growth of UPEC strains *in vitro*.

## Experimental procedures

### Bacterial growth conditions

All strains (Table 1) were cultured in lysogeny broth (LB) overnight at 37 °C, 200 rpm. UPEC strains CFT073 and UTI89 were isolated from the blood of patients with pyelonephritis (Mobley et al., 1990) and cystitis (Hultgren et al., 1986), respectively. The EC0 and EC1 bacteraemia isolates were a gift from Professor Thomas Evans’ laboratory and were isolated from the blood of hospitalised patients as described elsewhere (Goswami et al., 2018). For growth assays and growth for transcriptomic analysis, overnight cultures were diluted in culture medium (pooled human urine, MP-AU or enhanced AU), pre-warmed to 37 °C, to an OD _600 nm_ of 0.06. Cultures were incubated at 37 °C, 200 rpm for an 8 h period for growth assays, or until the desired OD _600 nm_ was reached for RNA extraction and subsequent transcriptomic analysis. For growth assays, the optical density of 1 mL neat samples was measured at hourly intervals.

**Table 1.** Strains used in this study.

| Name | Description | Phylogroup | Serotype | Reference |
| --- | --- | --- | --- | --- |
| CFT073 | Uropathogenic *E. coli* strain CFT073 | B2 | O6:H1 | (Mobley et al., 1990) |
| UTI89 | Uropathogenic *E. coli* strain UTI89 |  | O18:H7 | (Hultgren et al., 1986) |
| EC0_22 | *E. coli* isolate EC0_22 |  | O6:H1 | (Goswami et al., 2018) |
| EC0_32 | *E. coli* isolate EC0_32 |  | O1:H7 |  |
| EC0_38 | *E. coli* isolate EC0_38 |  | O25:H4 |  |
| EC0_58 | *E. coli* isolate EC0_58 |  | O18:H7 |  |
| EC0_62 | *E. coli* isolate EC0_62 |  | O2:H7 |  |
| EC1_57 | *E. coli* isolate EC1_57 |  | O18:H31 |  |
| EC1_60 | *E. coli* isolate EC1_60 |  | O22:H1 |  |
| EC1_77 | *E. coli* isolate EC1_77 |  | O16:H5 |  |
| EC1_84 | *E. coli* isolate EC1_84 |  | O4:H1 |  |
| EC1_91 | *E. coli* isolate EC1_91 |  | O18:H1 |  |
| EC1_93 | *E. coli* isolate EC1_93 |  | O6:H31 |  |

### Pooled human urine

The first morning urine from five healthy volunteers was pooled and filtered using a 0.22 μm vacuum filter (Sigma). Here, healthy describes the absence of infection. Volunteers were not asked to disclose other factors such as gender, diet, medication, or menstruation status. Pooled human urine was stored at 4 °C for use within four days or aliquoted and stored at - 20 °C for later use. Frozen aliquots were thawed at 4 °C overnight and brought to room temperature before use.

### Multipurpose artificial urine

MP-AU was adapted from the literature (Sarigul et al., 2019). All components except uric acid were prepared as 1 M or 0.5 M stock solutions in distilled deionised water and filter sterilised (Table 2). Uric acid was added in powder form as it is not easily dissolved in water. 1 M urea was prepared on the day of use. In general, MP-AU without urea was prepared the day before use.

**Table 2.** MP-AU components. The desired volume was made up with distilled deionised water.

| Component | Concentration in MP-AU | Supplier |
| --- | --- | --- |
| Sodium sulfate decahydrate | 11.965 mM | Sigma-Aldrich |
| Trisodium citrate | 2.450 mM | Sigma-Aldrich |
| Creatinine | 7.791 mM | Sigma-Aldrich |
| Urea | 249.750 mM | Fisher Scientific |
| Uric acid | 0.25 g/L | Sigma-Aldrich |
| Potassium chloride | 30.953 mM | Sigma-Aldrich |
| Sodium chloride | 30.953 mM | Sigma-Aldrich |
| Calcium chloride | 1.663 mM | Sigma-Aldrich |
| Ammonium chloride | 23.667 mM | Sigma-Aldrich |
| Potassium oxalate monohydrate | 0.190 mM | Sigma-Aldrich |
| Magnesium sulfate heptahydrate | 4.389 mM | Sigma-Aldrich |
| Sodium phosphate monobasic dihydrate | 18.667 mM | Sigma-Aldrich |
| Sodium phosphate dibasic dihydrate | 4.667 mM | Sigma-Aldrich |

### Enhanced AU

Supplement solutions were prepared in water and filter sterilised, with the exception of L-tyrosine, which was mixed in water but not filtered, due to its insolubility in water (Table S1). Supplements were added to MP-AU, with urea, at a final concentration of 5 mM on the day of use.

### Transcriptomic analysis

To directly compare the transcriptomic profile of UPEC strain CFT073 in MP-AU and pooled human urine, CFT073 was cultured in triplicate in M9 minimal media for 4 h. Bacteria were pelleted by centrifugation and resuspended in MP-AU or pooled human urine. After 1 h culture in MP-AU or pooled human urine, samples were taken and normalised to OD _600 nm_ ∼0.6. To compare the transcriptomic profile of CFT073 and UTI89 in enhanced AU and pooled human urine, UPEC strains CFT073 and UTI89 were cultured in triplicate in enhanced AU or pooled human urine until bacteria had reached an OD _600 nm_ of >0.4 (mid-exponential phase). Cultures were normalised to OD _600 nm_ 1.0. For all transcriptomic analysis experiments, bacteria were centrifuged at maximum speed for 1 min. Bacteria were treated with RNAprotect Bacteria Reagent (Qiagen) as per manufacturer’s instructions and stored at −20°C until RNA extraction within four days. Total RNA was isolated as per manufacturer’s instructions using the PureLink RNA Mini Kit (Invitrogen). DNA was removed using TURBO DNase (Invitrogen). RNA was further extracted using phenol:chloroform:isoamyl alcohol (Invitrogen) and was precipitated using 100% ethanol (Fisher Scientific). The concentration and purity of the extracted RNA was quantified using a spectrophotometer (DeNovix). The integrity of RNA was assessed by running gel electrophoresis to examine 23S and 16S rRNA bands. The presence of residual or contaminant DNA was determined by polymerase chain reaction to amplify *fimA*. Samples were sent to Glasgow Polyomics for ribosomal depletion using a QIAseq FastSelect 5S/16S/23S Kit (Qiagen), cDNA library preparation using a TruSeq Stranded Total RNA Library Prep Gold (Illumina) and RNA sequencing to a depth of 10 million 75 bp or 100 bp single end reads using a Nextseq500 or NextSeq2000 sequencing system (Illumina). Data were analysed using CLC Genomics Workbench version 21.0.5 (Qiagen). Reads were mapped to the CFT073 and UTI89 reference genomes (NCBI accession numbers NC_004431.1 and NC_007946.1, respectively) using default parameters. Differential expression was calculating using the Differential Expression for RNA-seq tool in CLC Genomics Workbench. Genes were taken as significantly differentially expressed with a fold change of ≥1.5 or ≤-1.5 and a false-discovery rate (FDR)-corrected *p*-value of ≤0.05. Volcano plots were generated using Prism version 9.5.1 (GraphPad). Functional categories of the differentially expressed genes (DEGs) were assigned using the UniProtKB gene ontology (GO) biological process annotation in the first instance, or the GO molecular function where the GO biological process annotation was not available (Bateman et al., 2021). Raw sequencing data have been deposited in the European Nucleotide Archive under the project accession number PRJEB55151.

### Metabolomics analysis

Metabolites were extracted from pooled human urine or MP-AU using an extraction solvent of chloroform:methanol:water in a 1:3:1 v/v/v ratio. The components for the extraction solvent were all high-performance liquid chromatography (HPLC) grade and supplied by Fisher Scientific. Extracted metabolites were analysed by liquid-chromatography mass spectrometry (LC-MS) by Glasgow Polyomics using a Dionex UltiMate 3000 RSLC system and an Q Exactive Orbitrap (Thermo Fisher Scientific). Data were analysed using the Glasgow Polyomics Integrated Metabolomics Pipeline (PiMP) software (Gloaguen et al., 2017). Statistical analysis, the Venn diagram and bar charts were generated using Prism version 9.5.1 (GraphPad).

### Iron quantification

ICP-MS of acidified pooled human urine, MP-AU and enhanced AU containing final concentration 1 mM iron (II) sulfate heptahydrate (Sigma-Aldrich) was performed by Dr Lorna Eades at the University of Edinburgh using the 8900 Triple Quadrupole ICP-MS instrument (Agilent). The samples were acidified with trace metal grade nitric acid (Fisher Scientific) and analytical reagent grade hydrochloric acid S.G. 1.18 (Fisher Scientific).

## Results

### MP-AU does not support the growth of UPEC strains in vitro

To determine whether MP-AU is suitable for the culture of UPEC strains *in vitro*, we performed growth assays of prototypic UPEC strains CFT073 and UTI89 in this media. The growth rates of these strains in MP-AU were compared to those obtained for growth in pooled human urine. Sarigul’s MP-AU did not support the optimal growth of UPEC strains CFT073 or UTI89 (Fig. 1a). The optical density of CFT073 and UTI89 in MP-AU decreased in the first hour of growth, suggesting that there was some lysis of the bacterial cells. Sarigul *et al*. suggests adding peptone and yeast extract to MP-AU for bacterial growth studies (Sarigul et al., 2019), however, these were omitted here due to their absence in healthy human urine and subsequent unsuitability for gene expression studies versus pooled human urine. To allow for transcriptomic analysis of CFT073 in MP-AU or pooled human urine, bacteria were first cultured in M9 minimal media before being resuspended in either MP-AU or pooled human urine. There were vast differences in the transcriptomic profile following 1 h exposure to MP-AU or pooled human urine, with 3325 differentially expressed genes in MP-AU compared to pooled human urine (Fig. 1c). This corresponds to a shift in 66% of the CFT073 genome. 1625 of these genes were upregulated, while 1690 were downregulated. The ten most significantly upregulated genes include those involved in arginine catabolism and carboxylic acid catabolism (*astABCDE, fadB*), as well as *argT*, which encodes a lysine/arginine/ornithine ABC transporter substrate-binding protein, *C_RS19130* (previously *c4034* or *yhdW*), and *yhdX* and *ytfT*, which encode ABC transporter permeases. The ten most significantly downregulated genes include *glnA*, *cyuA*, *gcvT*, *gcvH*, *lacY*, *rplC*, *lacA*, *rplD*, *sitC* and *chuY*.

**Fig 1.**
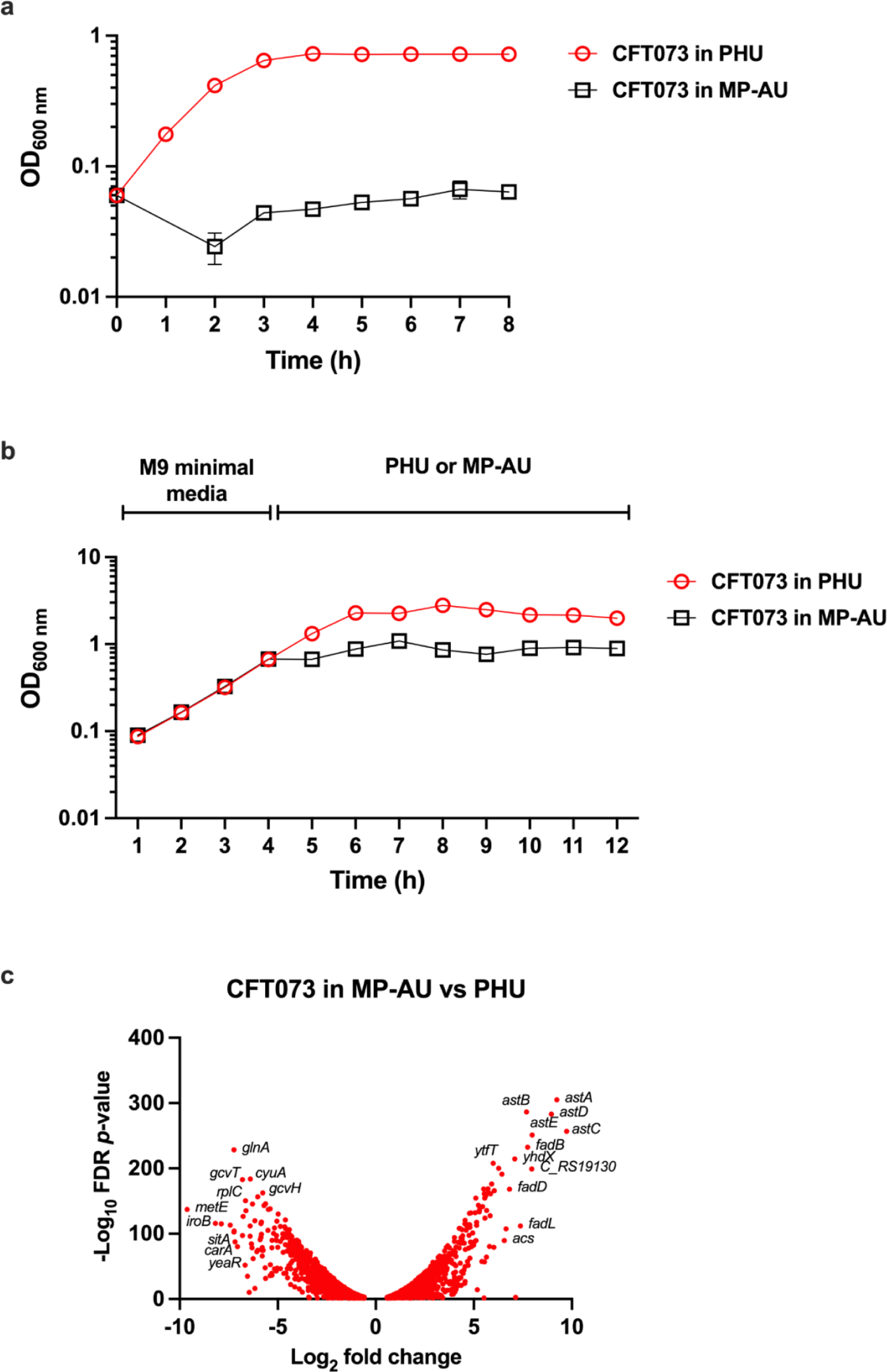
Assessing the suitability of multipurpose artificial urine for *in vitro* culture of uropathogenic *E. coli* strains. **a)** Overnight cultures of wild-type CFT073 were inoculated into pre-warmed filtered pooled human urine (PHU, red circles) or multipurpose artificial urine (MP-AU, black squares), at a starting OD _600 nm_ of 0.06. Error bars represent the standard error of the mean of three biological replicates. **b)** Overnight cultures of CFT073 were inoculated into pre-warmed M9 minimal media. After 4 h, cultures were pelleted by centrifugation and resuspended in PHU (red circles) or MP-AU (black squares). Three biological replicates were performed. **c)** Samples were removed following 1 h culture in PHU or MP-AU and normalised to OD _600 nm_ of 1 for RNA extraction. Volcano plot represents significant differentially expressed genes in MP-AU relative to PHU. FDR *p*-value ≤0.05, fold change ≥1.5 or ≤-1.5.

### Metabolomic analysis of pooled human urine

Sarigul *et al*. demonstrated that the levels of components in their MP-AU are similar healthy human urine (Sarigul et al., 2019). However, as the MP-AU did not facilitate the growth of UPEC strains *in vitro*, we hypothesised that there were components in healthy human urine that could be used to supplement MP-AU and support the growth of UPEC strains. LC-MS analysis was used to identify and compare the relative abundance of metabolites extracted from MP-AU and healthy human urine samples. Untargeted LC-MS identified 68 metabolites across all samples and putatively annotated a further 5703 metabolites, however as many of these shared a peak ID and could not be further distinguished by LC-MS, there were 65 true identified metabolites and 1289 true annotated metabolites (Table S2). 53 identified metabolites were present in both human urine and MP-AU, while 10 metabolites were only present in human urine, and two were only present in MP-AU (Fig. 2a). Of the 53 identified metabolites in common between the two media, only orthophosphate and nicotinamide were more abundant in MP-AU compared to human urine. The difference was statistically significant for orthophosphate (*p* value ≤0.05) but not for nicotinamide. The remaining 51 metabolites were more abundant in human urine. There was no statistical significance in the different average abundances of creatinine or oxalate between human urine and MP-AU. The increased abundance in human urine compared to MP-AU was statistically significant (*p* value ≤0.05) for six metabolites, highly significant (*p* value ≤0.01) for 26 metabolites and very highly significant (*p* value ≤0.001) for 18 metabolites. We chose to focus only on the identified metabolites, rather than annotated metabolites. The ratio of metabolites in MP-AU relative to human urine is shown in Fig. 2b.

**Fig 2.**
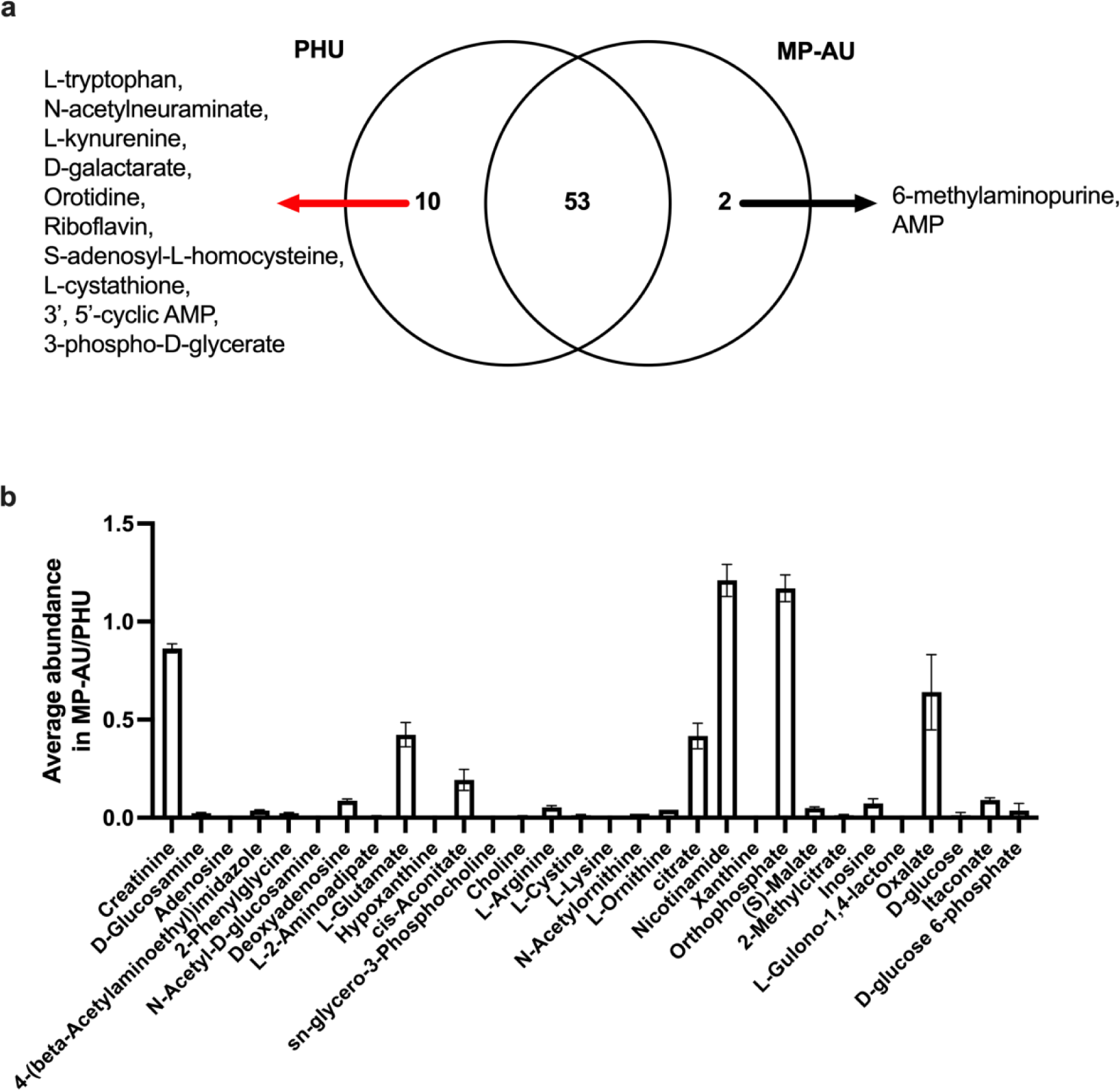
Determining the metabolites present in pooled human urine and multipurpose artificial urine by liquid chromatography mass spectrometrya) Venn diagram displaying the distribution of identified metabolites between pooled human urine (PHU) and multi-purpose artificial urine (MP-AU). **b**) The ratio of average abundance of identified metabolites in MP-AU relative to PHU. Metabolites with a ratio of average abundance of zero are excluded. Error bars represent the SEM of three technical replicates.

### Modifying MP-AU

Metabolites were added to MP-AU based on their relative abundance in pooled human urine samples over MP-AU, or studies published in the literature. Where LC-MS identified metabolites that were the product of another metabolite’s degradation or synthesis the by product was not included in the enhanced artificial urine formulation. For example, LC-MS analysis showed that L-kynurenine is present in pooled human urine. L-kynurenine is a by-product of tryptophan catabolism (van der Leek et al., 2017) and therefore L-tryptophan was used as a supplement instead of L-kynurenine. The majority of amino acids added were L-enantiomers, due to their physiological relevance in protein synthesis (Suzuki et al., 2021). There have also been reports of endogenous D-amino acids eukaryotes, such as D-glutamate (Katane et al., 2020), D-cysteine (Roychaudhuri and Snyder, 2022) and D-serine (Anfora et al., 2008; Cava et al., 2011), hence these were also added. While not in our LC-MS dataset, D-sorbitol is listed on the Urine Metabolome Database with detected quantities in the urine of adults at 3.5-9.9 μmol/mmol creatinine (Bouatra et al., 2013) and was included as it is suggested to be an important carbon source for UPEC strains in the urinary tract (Mann et al., 2017). The components were added to MP-AU at a final concentration of 0.5 M, and this was sufficient to support the growth of CFT073 and UTI89 (Fig. 3a). Both strains achieved the same OD _600 nm_ at 8 h growth in enhanced AU and the OD _600 nm_ of CFT073 was the same as that in pooled human urine. Iron is an important metal for the survival of UPEC strains in the urinary tract, with UPEC strains encoding a number of proteins that acquire, scavenge, increase the uptake of iron from urine. Sarigul’s MP-AU does not include iron but Brooks and Keevil’s artificial urine formulation contains small amounts of iron (II) sulfate (Brooks and Keevil, 1997; Sarigul et al., 2019). To assess iron levels in pooled human urine, we used inductively coupled plasma mass spectrometry (ICP-MS) to compare iron levels in pooled human urine with enhanced AU with supplemented with iron (II) sulfate at 1 mM. ICP-MS revealed that the levels of iron in human urine was negligible, while the levels in our enhanced artificial urine were sixty-fold higher (Fig. S1). As a result, iron (II) sulfate was omitted from subsequent formulations.

**Fig 3.**
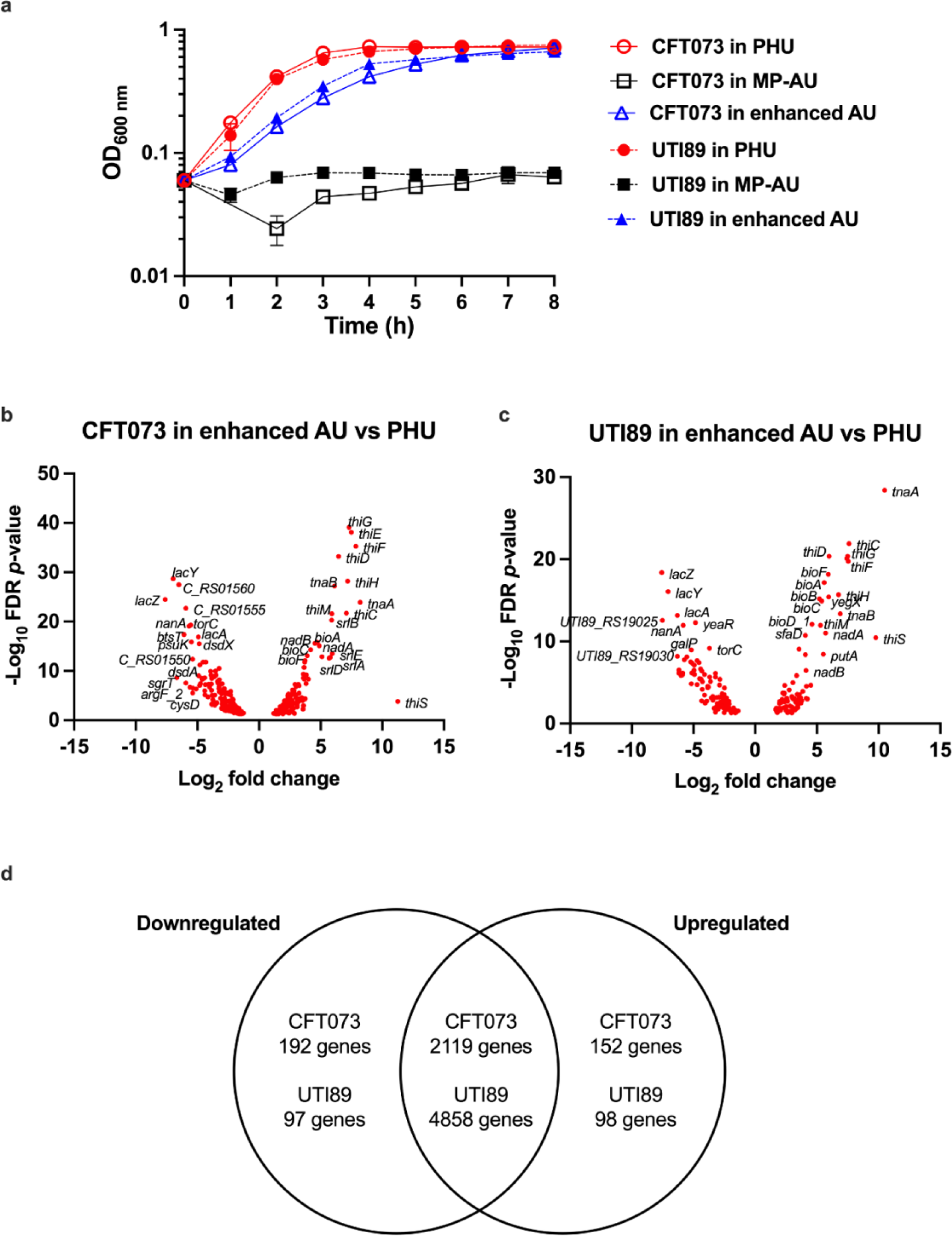
Growth in enhanced AU vs PHU and MP-AU. Overnight cultures of wild-type CFT073 or UTI89 were inoculated into pre-warmed filtered pooled human urine (PHU, red circles), multipurpose artificial urine (MP-AU, black squares) or enhanced artificial urine (enhanced AU, blue triangles) at a starting OD _600 nm_ of 0.06. Error bars represent the standard error of the mean of three biological replicates. **b)** Overnight cultures of CFT073 or UTI89 were inoculated into pre-warmed enhanced AU or PHU. Samples were removed during mid-exponential phase and normalised to OD _600 nm_ of 1 for RNA extraction. Volcano plot represents significant differentially expressed genes in enhanced AU relative to PHU. FDR *p*-value ≤0.05, fold change ≥1.5 or ≤-1.5. Three biological replicates were performed.

### Transcriptomic analysis in EnAU

Transcriptomics provides a global insight into how the bacterial cell responds to its environment. Subtle changes in the nutrient content of the growth media are reflected in distinct patterns of gene expression. Moreover, if enhanced AU is to be used to study UPEC *in vitro*, it is important to understand if key virulence genes are expressed to the same extent. To determine whether this was the case, transcriptomic analysis was performed on RNA extracted from CFT073 and UTI89 cultured in enhanced AU or pooled human urine. Samples for RNA extraction were taken during the mid-exponential phase to better capture the transcriptomic profile during growth in either condition. In contrast to the original MP-AU formulation where thousands of genes were affected, in this enhanced AU, 6.8% of the CFT073 genome (344 genes) and 3.8% of the UTI89 genome (189 genes) were significantly differentially expressed compared to pooled human urine (Fig. 3d). There were 152 significant upregulated genes in CFT073 in enhanced AU compared to pooled human urine, and 192 significant downregulated genes (Fig. 3b). In UTI89, there were 91 upregulated and 98 downregulated genes, including the hypothetical protein-encoding gene *UTI89_RS29285* carried on the UTI89 plasmid (Fig. 3c). The 50 most significant differentially expressed genes in CFT073 and UTI89 are shown in Tables S3 and S4, respectively.

### Expression of virulence factors in EnAU

Promisingly, a number of genes associated with virulence were similarly expressed (not significantly up or downregulated) in CFT073 or UTI89 in EnAU and PHU. We examined genes known to be associated with virulence, such as those encoding toxins, adhesins, siderophores and iron uptake systems.

Almost half of all UPEC strains secrete the pore-forming toxin hemolysin (Nhu et al., 2019). Hemolysin is encoded for by *hlyA*, which is similarly expressed in CFT073 and UTI89 in EnAU and pooled human urine. Another toxin, cytotoxic necrotising factor 1 or CNF1, is an important toxin expressed by some UPEC strains, including UTI89, but it is not encoded for by the CFT073 genome (Smith et al., 2008). Here, there was a similar level of expression of *cnf1* in UTI89 in EnAU compared to PHU. CFT073, but not UTI89, produces the serine protease autotransporter toxins Pic and Sat. These genes (*pic* and *sat,* respectively) are similarly expressed in both EnAU and in PHU. The colibactin genotoxin is expressed during UTI in humans, and is associated with DNA damage in the urothelial cells in a mouse model of UTI (Chagneau et al., 2021). 17 CFT073 *clb* genes and 15 UTI89 *clb* genes, which encode colibactin, were similarly expressed in EnAU and pooled human urine.

Adherence to the uroepithelium through the expression of adhesins is a critical step in uropathogenesis. Six genes in the type 1 fimbriae operon (*fimC*, *fimD, fimF, fimG, fimH* and *fimI*) were significantly upregulated in CFT073 in EnAU compared to in PHU. In contrast, only three type 1 fimbriae genes (*fimF, fimG* and *fimH*) in UTI89 were significantly upregulated in EnAU compared to PHU. The adhesin FimH was significantly upregulated three-fold in CFT073 and 3.5-fold in UTI89 cultured in EnAU compared with PHU.

Perhaps this is unsurprising, as Greene *et al*. found that culture in filtered human urine induces the phase OFF phase of UTI89 *fimS* (Greene et al., 2015). While *fim* operon expression is upregulated in EnAU compared to in pooled human urine, this is still an advantage for the further study of type 1 fimbriae expression and function under different conditions. The pyelonephritis-associated pili (pap) or P fimbriae, thought to be associated with adhesion in the kidney, are similarly expressed in CFT073 in EnAU and PHU. Of all the *pap* operon genes in CFT073, only *papH_1* is significantly differentially expressed in EnAU relative to pooled human urine (2.6-fold increase). *CsgBA* are the minor and major subunits of the curli fibres associated with adhesion and are similarly expressed in CFT073 in EnAU and in PHU, however c*sgC* is significantly downregulated. In UTI89, the *csgBAC* operon is similarly expressed in the two media. The S and F1C fimbriae are encoded for by the *sfa* and *foc* genes, which are also similarly expressed by CFT073 in EnAU and in PHU.

Survival and growth in the urinary tract requires the uptake of iron from a nutritionally-poor environment via iron-uptake systems and siderophores that are secreted by UPEC strains (Frick-Cheng et al., 2022). The *sitABCD* genes, which encode an iron/manganese ABC transporter, are similarly expressed in CFT073 and UTI89 in EnAU compared to pooled human urine. Genes for TonB-dependent heme receptors ChuA (*chuA*) and Hma (*C_RS11765, UTI89_C2234*) are also similarly expressed by CFT073 and UTI89 in both media.

Siderophores aerobactin, enterobactin, salmochelin and yersiniabactin are encoded by UPEC strains to bind to iron sequestered by the host, for example in haemoglobin (Frick-Cheng et al., 2022). Aerobactin genes *iucABCD* are similarly expressed in CFT073 in EnAU and in PHU, so too is the gene encoding its receptor *iutA*. In CFT073 and UTI89, enterobactin genes *entCEBAH*, *entD* and *entS*, *fepA* (encoding the enterobactin receptor), salmochelin genes *iroBCDE* and *iroN* (encoding the salmochelin receptor) were similarly expressed in EnAU and pooled human urine. Of the yersiniabactin genes, only *ybtX*, which encodes the yersiniabactin-associated zinc MSF transporter YbtX, is significantly upregulated in CFT073 in EnAU compared to in pooled human urine. The other yersiniabactin genes are similarly expressed. None of the UTI89 *ybt* genes were significantly differentially expressed in EnAU compared to PHU.

### Suitability of enhanced AU to study other UPEC strains

While CFT073 and UTI89 are prototypic model strains for UPEC study, there is great diversity in the range of isolates associated with UTIs. A versatile artificial urine must be robust enough to support the growth of a variety of strains. To determine the usefulness of our enhanced artificial urine, growth assays were performed on isolates taken from patients who had been catheterised, and subsequently developed blood infections. The isolates were varied in serotype but all from the B2 phylogroup that is strongly associated with UPEC infections. Enhanced AU supported the growth of these clinical isolates demonstrating that it is useful in supporting the growth of strains, and not just the prototypical laboratory strains (Fig. 4).

**Fig 4.**
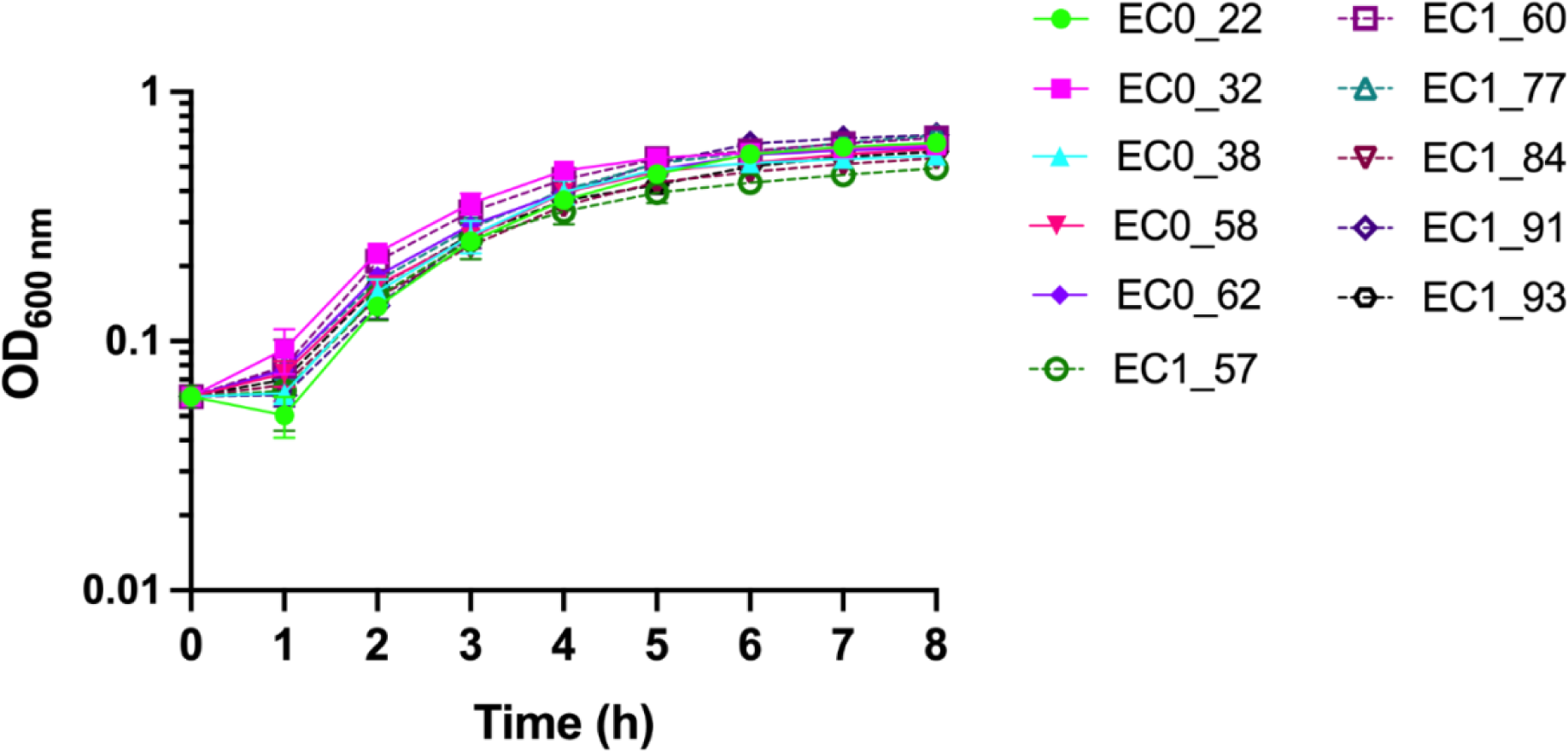
Growth in enhanced AU. Overnight cultures of bacteraemia isolates were inoculated into pre-warmed enhanced artificial urine at a starting OD _600 nm_ of 0.06. Error bars represent the standard error of the mean of three biological replicates.

## Discussion

Advances in formulation of artificial urine have meant that there are a number of different formulations proposed in the literature or available commercially. Some artificial urine formulations used to culture bacterial strains *in vitro* contain components not normally present in healthy human urine. Here, we attempted to modify a multi-purpose artificial urine to better resemble pooled human urine in terms of the ability to support growth of UPEC strains and to induce a similar transcriptomic profile during growth. A multipurpose artificial urine described in the literature was not suitable for the culture of UPEC strains CFT073 and UTI89 when peptone and yeast extract were omitted. Peptone and yeast extract used in some artificial urine formulations act as sources of amino acids, short peptides, trace elements and nucleic acids that may be normally present in human urine (Brooks and Keevil, 1997). As the quantities of components in commercially available reagents such as peptone and yeast extract are not always available, our approach of supplementing MP-AU with specific quantities of amino acids and metabolites aims to generate a more defined and reproducible medium. The complexity of urine suggests that there are certain factors in the urine that allow for growth and survival of those UPEC strains *in vitro*. As revealed by LC-MS, there were 65 identified metabolites that varied in abundance between pooled human urine and MP-AU. Unfortunately, LC-MS can indicate relative abundances but cannot be used to precisely quantify the metabolites present in a sample. Additionally, LC-MS cannot identify all of the metabolites present in a sample. LC-MS was used here only to identify metabolites in pooled human urine that could be used to supplement AU. It is worth noting that these studies were all performed on filtered urine. Filter sterilisation may result in the loss of components that contribute to bacterial growth and/or gene expression *in vivo*. Furthermore, *in vivo*, there is constant voiding and replenishing of urine in the bladder, whereas in our *in vitro* experiments, the media is not replenished. Nevertheless, similar levels of transcription between the two media suggests that EnAU does can in fact mimic conditions of PHU and we have generated an artificial urine that is consistent in allowing for growth of multiple UPEC strains for studying expression of virulence genes *in vitro*.

## Supporting information

Supplementary files complete

## Data availability statement

Raw sequencing data have been deposited in the European Nucleotide Archive under the project accession number PRJEB55151.

## Acknowledgements

The authors thank Glasgow Polyomics, Dr Lorna Eades at the University of Edinburgh, and Professor Thomas Evans, Dr Cosmika Goswami and Dr Stephen Fox at the University of Glasgow for their help in this work. For the purpose of open access, the authors have applied a Creative Commons Attribution (CC BY) licence to any Author Accepted Manuscript version arising from this submission.

