## Supplementary files complete for "Enhancing a multipurpose artificial urine for culture and gene expression studies of uropathogenic *Escherichia coli* strains"

**This PDF Includes:**

Tables S1-S4

Figure S1

**Table S1.** Supplements added to MP-AU to prepare enhanced AU. Supplements were added at a final concentration of 0.5 mM.

| **Component** | **Supplier** |
|---|---|
| D-alanine | Sigma-Aldrich |
| D-serine | |
| D-sorbitol | |
| Glycine | |
| Hippuric acid | |
| L-alanine | |
| L-arginine | |
| L-aspartic acid | |
| L-carnitine | |
| L-citrulline | |
| L-cysteine hydrochloride | |
| L-glutamic acid monosodium salt hydrate | |
| L-glutamine | |
| L-histidine | |
| L-lysine | |
| L-methionine | |
| L-ornithine monohydrochloride | |
| L-proline | |
| L-serine | |
| L-tryptophan | |
| L-tyrosine | |

**Table S2.** Abundance of metabolites identified in pooled human urine vs multi-purpose artificial urine.

| Name | Formula | Log fold change |
|---|---|---|
| N-Acetylglutamine | $C_7H_{12}N_2O_4$ | 12.00 |
| L-Citrulline | $C_6H_{13}N_3O_3$ | 10.85 |
| L-Carnitine | $C_7H_{15}NO_3$ | 10.77 |
| N(pi)-Methyl-L-histidine | $C_7H_{11}N_3O_2$ | 10.68 |
| Glycine | $C_2H_5NO_2$ | 10.47 |
| L-Tryptophan | $C_{11}H_{12}N_2O_2$ | 10.03 |
| D-Galacturonate | $C_6H_{10}O_7$ | 9.66 |
| L-Glutamine | $C_5H_{10}N_2O_3$ | 9.36 |
| D-Threose | $C_4H_8O_4$ | 9.26 |
| L-Tyrosine | $C_9H_{11}NO_3$ | 9.24 |
| D-Ribose | $C_5H_{10}O_5$ | 9.20 |
| O-Acetylcarnitine | $C_9H_{17}NO_4$ | 9.16 |
| 2-Oxoglutarate | $C_5H_6O_5$ | 9.11 |
| Ala-Gly | $C_5H_{10}N_2O_3$ | 9.10 |
| sucrose | $C_{12}H_{22}O_{11}$ | 9.10 |
| Betaine | $C_5H_{11}NO_2$ | 8.56 |
| N2-Acetyl-L-lysine | $C_8H_{16}N_2O_3$ | 8.45 |
| L-Serine | $C_3H_7NO_3$ | 8.42 |
| N-Acetylneuraminate | $C_{11}H_{19}NO_9$ | 8.39 |
| Hypoxanthine | $C_5H_4N_4O$ | 8.39 |
| Pantothenate | $C_9H_{17}NO_5$ | 8.27 |
| D-Gluconic acid | $C_6H_{12}O_7$ | 8.25 |
| N-acetyl-L-glutamate | $C_7H_{11}NO_5$ | 8.24 |
| L-Aspartate | $C_4H_7NO_4$ | 8.22 |
| L-Methionine | $C_5H_{11}NO_2S$ | 8.00 |
| 5-Oxoproline | $C_5H_7NO_3$ | 7.86 |
| L-2-Aminoadipate | $C_6H_{11}NO_4$ | 7.55 |

| | | |
|---|---|---|
| L-Kynurenine | $C_{10}H_{12}N_2O_3$ | 7.52 |
| Choline | $C_5H_{13}NO$ | 7.46 |
| sn-glycero-3-Phosphocholine | $C_8H_{20}NO_6P$ | 7.27 |
| Xanthine | $C_5H_4N_4O_2$ | 7.19 |
| Adenosine | $C_{10}H_{13}N_5O_4$ | 7.18 |
| D-Galactarate | $C_6H_{10}O_8$ | 7.10 |
| L-Lysine | $C_6H_{14}N_2O_2$ | 7.09 |
| Orotidine | $C_{10}H_{12}N_2O_8$ | 6.94 |
| N-Acetyl-D-glucosamine | $C_8H_{15}NO_6$ | 6.94 |
| riboflavin | $C_{17}H_{20}N_4O_6$ | 6.86 |
| L-Gulono-1,4-lactone | $C_6H_{10}O_6$ | 6.69 |
| L-Cystine | $C_6H_{12}N_2O_4S_2$ | 6.50 |
| 2-Methylcitrate | $C_7H_{10}O_7$ | 6.50 |
| S-Adenosyl-L-homocysteine | $C_{14}H_{20}N_6O_5S$ | 6.49 |
| L-Cystathionine | $C_7H_{14}N_2O_4S$ | 5.98 |
| 3',5'-Cyclic AMP | $C_{10}H_{12}N_5O_6P$ | 5.97 |
| N-Acetylornithine | $C_7H_{14}N_2O_3$ | 5.95 |
| D-Glucosamine | $C_6H_{13}NO_5$ | 5.38 |
| 2-Phenylglycine | $C_8H_9NO_2$ | 5.34 |
| D-glucose | $C_6H_{12}O_6$ | 4.86 |
| 4-(beta-Acetylaminoethyl)imidazole | $C_7H_{11}N_3O$ | 4.79 |
| L-Ornithine | $C_5H_{12}N_2O_2$ | 4.70 |
| (S)-Malate | $C_4H_6O_5$ | 4.29 |
| L-Arginine | $C_6H_{14}N_4O_2$ | 4.23 |
| 3-Phospho-D-glycerate | $C_3H_7O_7P$ | 4.08 |
| Inosine | $C_{10}H_{12}N_4O_5$ | 3.89 |
| Deoxyadenosine | $C_{10}H_{13}N_5O_3$ | 3.49 |
| Itaconate | $C_5H_6O_4$ | 3.46 |
| D-glucose 6-phosphate | $C_6H_{13}O_9P$ | 3.37 |

| | | |
|---|---|---|
| cis-Aconitate | $C_6H_6O_6$ | 2.49 |
| citrate | $C_6H_8O_7$ | 1.30 |
| L-Glutamate | $C_5H_9NO_4$ | 1.27 |
| Oxalate | $C_2H_2O_4$ | 0.76 |
| Creatinine | $C_4H_7N_3O$ | 0.21 |
| Orthophosphate | $H_3O_4P$ | -0.22 |
| Nicotinamide | $C_6H_6N_2O$ | -0.27 |
| 6-Methylaminopurine | $C_6H_7N_5$ | -1.88 |
| AMP | $C_{10}H_{14}N_5O_7P$ | -2.25 |

**Table S3.** 50 most significant differentially expressed genes in CFT073 in enhanced AU vs pooled human urine.

| Name | Gene product | Function | Fold change | FDR p-value |
|---|---|---|---|---|
| *thiG* | thiazole synthase | Thiamine diphosphate biosynthetic process | 158.89 | 7.56E-40 |
| *thiE* | thiamine phosphate synthase | Thiamine biosynthetic process, thiamine diphosphate biosynthetic process | 179.03 | 7.70E-39 |
| *thiF* | thiazole biosynthesis adenylyltransferase ThiF | ATP binding, metal ion binding, nucleotidyltransferase activity, ubiquitin-like modifier activating enzyme activity | 230.97 | 4.64E-36 |
| *thiD* | bifunctional hydroxymethylpyrimidine kinase/phosphomethylpyrimidine kinase | Phosphorylation, thiamine biosynthetic process, thiamine diphosphate biosynthetic process | 87.54 | 5.85E-34 |
| *lacY* | lactose permease | Carbohydrate:proton symporter activity | -126.12 | 1.79E-29 |
| *thiH* | 2-iminoacetate synthase ThiH | Thiamine biosynthetic process | 146.37 | 6.42E-29 |
| *C_RS01560* | PTS sugar transporter subunit IIA | Not available | -90.41 | 2.84E-28 |
| *tnaB* | low affinity tryptophan permease TnaB | Aromatic amino acid transmembrane transporter activity | 69.48 | 5.66E-28 |
| *lacZ* | beta-galactosidase | Carbohydrate catabolic process | -196.98 | 3.18E-25 |
| *tnaA* | tryptophanase | Tryptophanase activity | 295.36 | 1.20E-24 |
| *C_RS01555* | PTS sugar transporter subunit IIB | Phosphoenolpyruvate-dependent sugar phosphotransferase system | -61.38 | 1.79E-23 |
| *thiC* | phosphomethylpyrimidine synthase ThiC | Thiamine biosynthetic process, thiamine diphosphate biosynthetic process | 135.21 | 1.93E-22 |
| *thiM* | hydroxyethylthiazole kinase | Phosphorylation, thiamine biosynthetic process, thiamine diphosphate biosynthetic process | 59.69 | 2.36E-22 |
| *srlB* | PTS glucitol/sorbitol transporter subunit IIA | Phosphoenolpyruvate-dependent sugar phosphotransferase system | 58.84 | 5.14E-21 |
| *torC* | pentaheme c-type cytochrome TorC | Anaerobic respiration | -46.31 | 4.72E-20 |
| *nanA* | N-acetylneuraminate lyase | Carbohydrate metabolic process, N-acetylneuraminate catabolic process | -51.54 | 7.49E-20 |

| | | | | |
|---|---|---|---|---|
| *btsT* | pyruvate/proton symporter BtsT | Cellular response to starvation | -68.15 | 4.38E-18 |
| *lacA* | galactoside O-acetyltransferase | Acetyltransferase activity | -30.13 | 1.25E-17 |
| *psuK* | pseudouridine kinase | Phosphorylation | -44.63 | 1.25E-16 |
| *bioA* | adenosylmethionine--8-amino-7-oxononanoate transaminase | Biotin biosynthetic process | 26.59 | 1.68E-16 |
| *nadB* | L-aspartate oxidase | NAD biosynthetic process | 23.09 | 2.38E-16 |
| *dsdX* | D-serine transporter DsdX | D-serine transport | -28.73 | 3.08E-16 |
| *nadA* | quinolinate synthase NadA | NAD biosynthetic process | 29.70 | 8.49E-16 |
| *bioC* | malonyl-ACP O-methyltransferase BioC | Biotin biosynthetic process, methylation | 18.26 | 4.64E-15 |
| *srlE* | PTS glucitol/sorbitol transporter subunit IIB | Phosphoenolpyruvate-dependent sugar phosphotransferase system | 60.96 | 3.00E-14 |
| *bioF* | 8-amino-7-oxononanoate synthase | Biotin biosynthetic process | 14.90 | 7.64E-14 |
| *gutM* | transcriptional regulator GutM | Not available | 34.26 | 1.24E-13 |
| *srlA* | PTS glucitol/sorbitol transporter subunit IIC | Phosphoenolpyruvate-dependent sugar phosphotransferase system | 54.82 | 1.86E-13 |
| *srlD* | sorbitol-6-phosphate dehydrogenase | Sorbitol-6-phosphate 2-dehydrogenase activity | 51.28 | 2.55E-13 |
| *C_RS01550* | PTS ascorbate transporter subunit IIC | Phosphoenolpyruvate-dependent sugar phosphotransferase system | -41.66 | 3.89E-13 |
| *bioB* | biotin synthase BioB | Biotin biosynthetic process | 13.34 | 7.42E-13 |
| *hiuH* | hydroxyisourate hydrolase | Purine nucleobase metabolic process | -19.97 | 1.65E-12 |
| *C_RS19900* | hypothetical protein | Not available | -23.19 | 1.65E-12 |
| *bioD_1* | dethiobiotin synthase | Biotin biosynthetic process | 13.03 | 1.65E-12 |
| *dsdA* | D-serine ammonia-lyase | D-amino acid metabolic process | -27.81 | 5.91E-12 |
| *zraP* | zinc resistance sensor/chaperone ZraP | Not available | 12.62 | 2.08E-11 |
| *C_RS11805* | DeoR family transcriptional regulator | D-ribose catabolic process | -9.40 | 3.46E-11 |
| *nanE* | N-acetylmannosamine-6-phosphate 2-epimerase | Carbohydrate metabolic process, N-acetylmannosamine metabolic process, N-acetylneuraminate catabolic process | -11.48 | 1.03E-10 |

| | | | | |
|---|---|---|---|---|
| *uxaC* | glucuronate isomerase | Glucuronate catabolic process | -11.71 | 1.04E-10 |
| *ygbJ* | NAD(P)-dependent oxidoreductase | Carbohydrate metabolic process, organic acid catabolic process | -15.30 | 1.13E-10 |
| *yegX* | glycoside hydrolase family 25 protein | Cell wall macromolecule catabolic process, peptidoglycan catabolic process | 12.10 | 4.51E-10 |
| *araD_2* | L-ribulose-5-phosphate 4-epimerase | L-arabinose catabolic process to xylulose 5-phosphate | 14.27 | 6.66E-10 |
| *galP* | galactose/proton symporter | Carbohydrate transport | -11.23 | 7.08E-10 |
| *glpD* | glycerol-3-phosphate dehydrogenase | Glycerol-3-phosphate metabolic process | -15.03 | 7.08E-10 |
| *psuG* | pseudouridine-5'-phosphate glycosidase | Not available | -29.64 | 9.87E-10 |
| *nanT* | sialic acid transporter NanT | Carbohydrate transport | -13.07 | 1.12E-09 |
| *nanK* | N-acetylmannosamine kinase | N-acetylmannosamine metabolic process, N-acetylneuraminate catabolic process | -12.39 | 1.62E-09 |
| *sgrT* | glucose uptake inhibitor SgrT | Not available | -101.52 | 2.28E-09 |
| *idnD* | L-idonate 5-dehydrogenase | L-idonate 5-dehydrogenase activity | -21.06 | 2.55E-09 |
| *astE* | succinylglutamate desuccinylase | Arginine catabolic process to glutamate, arginine catabolic process to succinate | 12.17 | 2.83E-09 |

**Table S4.** 50 most significant differentially expressed genes in UTI89 in enhanced AU vs pooled human urine.

| **Name** | **Gene product** | **Functional group** | **Fold change** | **FDR *p*-value** |
|---|---|---|---|---|
| *tnaA* | tryptophanase | Tryptophanase activity | 1446.32 | 3.79E-29 |
| *thiC* | phosphomethylpyrimidine synthase ThiC | Thiamine biosynthetic process, thiamine diphosphate biosynthetic process | 196.09 | 1.19E-22 |
| *thiD* | bifunctional hydroxymethylpyrimidine kinase/phosphomethylpyrimidine kinase | Phosphorylation, thiamine biosynthetic process, thiamine diphosphate biosynthetic process | 63.66 | 4.39E-21 |
| *thiG* | thiazole synthase | Thiamine diphosphate biosynthetic process | 179.56 | 4.39E-21 |
| *thiE* | thiamine phosphate synthase | Thiamine biosynthetic process, thiamine diphosphate biosynthetic process | 171.08 | 7.93E-21 |
| *thiF* | thiazole biosynthesis adenylyltransferase ThiF | ATP binding, metal ion binding, nucleotidyltransferase activity, ubiquitin-like modifier activating enzyme activity | 187.10 | 1.76E-20 |
| *lacZ* | beta-galactosidase | Carbohydrate catabolic process | -191.23 | 4.11E-19 |
| *bioF* | 8-amino-7-oxononanoate synthase | Biotin biosynthetic process | 60.96 | 6.99E-19 |
| *bioA* | adenosylmethionine--8-amino-7-oxononanoate transaminase | Biotin biosynthetic process | 48.36 | 6.73E-18 |
| *lacY* | lactose permease | Carbohydrate:proton symporter activity | -132.55 | 8.65E-17 |
| *thiH* | 2-iminoacetate synthase ThiH | Thiamine biosynthetic process | 107.68 | 2.05E-16 |
| *yegX* | glycoside hydrolase family 25 protein | Cell wall macromolecule catabolic process, peptidoglycan catabolic process | 62.45 | 3.98E-16 |
| *bioB* | biotin synthase BioB | Biotin biosynthetic process | 36.95 | 6.57E-16 |
| *bioC* | malonyl-ACP O-methyltransferase BioC | Biotin biosynthetic process, methylation | 41.93 | 1.20E-15 |
| *tnaB* | low affinity tryptophan permease TnaB | Aromatic amino acid transmembrane transporter activity | 119.04 | 4.39E-14 |
| *lacA* | galactoside O-acetyltransferase | Acetyltransferase activity | -79.66 | 6.91E-14 |
| *UTI89_RS19025* | hypothetical protein | Not available | -185.23 | 2.75E-13 |

| | | | | |
|---|---|---|---|---|
| *yeaR* | DUF1971 domain-containing protein YeaR | Not available | -28.61 | 5.16E-13 |
| *bioD_1* | dethiobiotin synthase | Biotin biosynthetic process | 24.26 | 7.99E-13 |
| *nanA* | N-acetylneuraminate lyase | Carbohydrate metabolic process, N-acetylneuraminate catabolic process | -57.06 | 1.12E-12 |
| *thiM* | hydroxyethylthiazole kinase | Thiamine biosynthetic process, thiamine diphosphate biosynthetic process | 39.03 | 1.12E-12 |
| *nadA* | quinolinate synthase NadA | NAD biosynthetic process | 52.50 | 9.70E-12 |
| *sfaD* | S-fimbrial adhesin minor subunit SfaS | Cell adhesion | 16.80 | 1.87E-11 |
| *thiS* | sulfur carrier protein ThiS | Thiamine biosynthetic process | 879.87 | 3.46E-11 |
| *torC* | pentaheme c-type cytochrome TorC | Anaerobic respiration | -13.11 | 6.82E-10 |
| *UTI89_RS05345* | fimbria/pilus periplasmic chaperone | Regulation of DNA-templated transcription | 11.76 | 8.86E-10 |
| *galP* | galactose/proton symporter | Transmembrane transporter activity | -36.07 | 1.06E-09 |
| *putA* | trifunctional transcriptional regulator/proline dehydrogenase/L-glutamate gamma-semialdehyde dehydrogenase | Proline biosynthetic process, proline catabolic process to glutamate | 45.79 | 3.56E-09 |
| *UTI89_RS05360* | fimbrial protein | Cell wall organisation, chaperone-mediated protein folding | 16.87 | 3.99E-09 |
| *UTI89_RS19030* | hypothetical protein | Not available | -79.50 | 6.63E-09 |
| *uidB* | glucuronide transporter | Carbohydrate transport, sodium ion transport | -46.55 | 7.75E-09 |
| *uidA* | beta-glucuronidase | Carbohydrate catabolic process | -53.60 | 1.68E-08 |
| *yoaG* | DUF1869 domain-containing protein | Not available | -26.05 | 2.13E-08 |
| *UTI89_RS24120* | pyridoxal phosphate-dependent aminotransferase | Biosynthetic process | -37.27 | 3.00E-08 |
| *nanE* | N-acetylmannosamine-6-phosphate 2-epimerase | Carbohydrate metabolic process, N-acetylneuraminate metabolic process, N-acetylneuraminate catabolic process | -23.45 | 3.78E-08 |
| *ygjI* | amino acid permease | Transmembrane transporter activity | -34.88 | 4.27E-08 |

| | | | | |
|---|---|---|---|---|
| *ebgC* | beta-galactosidase subunit beta | Metabolic process | -30.65 | 7.74E-08 |
| *nanC* | N-acetylneuraminic acid outer membrane channel NanC | Carbohydrate transport | -21.57 | 2.35E-07 |
| *ygbL* | aldolase | Monosaccharide metabolic process | -72.54 | 3.12E-07 |
| *idnO* | gluconate 5-dehydrogenase | Gluconate 5-dehydrogranase activity | -31.18 | 3.27E-07 |
| *nadB* | L-aspartate oxidase | NAD biosynthetic process | 17.51 | 3.47E-07 |
| *glpB* | glycerol-3-phosphate dehydrogenase subunit GlpB | Glycerol catabolic process, glycerol-3-phosphate metabolic process | -20.08 | 4.20E-07 |
| *btsT* | pyruvate/proton symporter BtsT | Cellular response to starvation | -35.12 | 6.90E-07 |
| *idnD* | L-idonate 5-dehydrogenase | L-idonate 5-dehydrogenase activity | -33.74 | 7.16E-07 |
| *ygbK* | four-carbon acid sugar kinase family protein | Carbohydrate metabolic process, phosphorylation | -58.36 | 7.16E-07 |
| *ygbN* | GntP family transporter | Methylation | -72.32 | 7.16E-07 |
| *UTI89_RS22505* | CidA/LrgA family protein | Not available | -9.36 | 7.79E-07 |
| *glpD* | glycerol-3-phosphate dehydrogenase | Glycerol-3-phosphate metabolic process | -30.35 | 1.08E-06 |
| *nanK* | N-acetylmannosamine kinase | N-acetylmannoseamine metabolic process, N-acetylneuraminate catabolic process | -17.25 | 1.08E-06 |
| *ygbJ* | NAD(P)-dependent oxidoreductase | Organic acid catabolic process | -60.38 | 1.20E-06 |

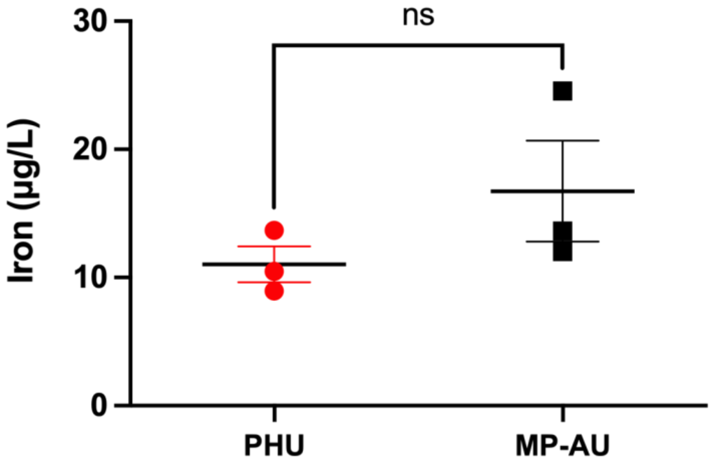


**Fig S1.** Determining the iron concentration in pooled human urine by inductively-coupled plasma mass spectrometry. The concentration of iron in PHU (red circles) and MP-AU (black squares) was analysed by ICP-MS. Data represent the mean and standard error of the mean of three technical replicates. Statistical significance was determined using Welch's t test.
